# Climate gradients drive the evolution of seed morphology and life history with impacts to seedling fitness in *Fraxinus nigra*

**DOI:** 10.64898/2026.04.17.719241

**Authors:** Kyra R. LoPiccolo, Jacob D. Mazza, Lucia A. Anderson, Dolzodmaa Davaasuren, Jill A. Hamilton

## Abstract

**Background and Aims:** Climate gradients influence seed morphology, emergence, and early life-history traits with cumulative impacts to individual fitness. For *ex situ* seed collections, which represent an invaluable repository of potential trait information for species management and conservation, climate data can guide preservation of adaptive variation and inform deployment strategies for restoration. Here we leverage a range-wide *ex situ* seed collection of critically endangered black ash seeds (*Fraxinus nigra*) to evaluate how climatic gradients shape variation in morphology and early life-history.

**Methods:** To test how climate of origin, seed morphology, and early life-history interact to impact first year fitness, high-throughput X-ray imaging and neural network–based segmentation were used to quantify variation in seed morphology for 701 maternal lineages spanning 76 populations across the range of *F. nigra*. Following this, a subset of seeds were used to establish a common garden experiment and quantify variation in emergence, early life-history transitions, and their cumulative impact to first-year survival and growth.

**Results:** On average, differences within-population explained ∼43% of the variability in seed morphology, while among-population differences explained ∼14%. This suggests that substantial genetic variation exists within populations for natural selection to act upon and differences have evolved among populations. Climate associations indicated warmer and drier environments predicted heavier seeds with faster developmental transitions and increased first-year height. Together, climate of origin, seed mass, and timing of developmental transitions best predicted cumulative fitness, with populations from more continental environments exhibiting greater survival and first-year height accumulation on average.

**Conclusions:** Overall, these results highlight the importance of climate of origin, seed traits, and early developmental transitions to first-year fitness in a perennial tree species. This work demonstrates how *ex situ* collections can be used to identify climatically structured trait variation and guide conservation strategies aimed at maintaining adaptive potential under environmental change.

## Introduction

Climate gradients shape the evolution of seed traits, with important consequences to population establishment and persistence under global change (Donohue *et al*., 2010; Cochrane *et al*., 2015; Saatkamp *et al*., 2019). Traits such as seed size, weight, and length vary predictably with temperature and precipitation, suggesting adaptation to local climatic conditions (Wright and Westoby, 1999; Moles and Westoby, 2003; Moles *et al*., 2005; Liu *et al*., 2013). These traits influence key components of fitness, including emergence timing (Seiwa, 2000; Verdú and Traveset, 2005), early growth (Castro, 1999; Baraloto *et al*., 2005; Carles *et al*., 2009), and survival (Moles and Westoby, 2004 *a*), with cumulative impacts to overall plant fitness (Sage *et al*., 2011; Center *et al*., 2016; Warwell and Shaw, 2019). However, given the pace of climate change, historic climate-trait relationships may be decoupled with impacts to population fitness (Howe *et al*., 2003; Savolainen *et al*., 2007; Hereford, 2009; Sáenz-Romero *et al*., 2016). Therefore, quantifying how climate, seed, and early-life history traits influence fitness or proxies thereof is needed to predict population responses to global change (Cochrane *et al*., 2015). For *ex situ* seed collections, these insights can guide supplemental collection priorities, ensure maintenance of adaptive variation, and inform seed deployment strategies for restoration in a changing climate.

Seed trait variability can reflect trade-offs associated with varying selective pressures that differ across environments. Heritable traits, including seed size and weight are generally greater in warmer and drier environments (Leishman *et al*., 2000; Moles *et al*., 2005; Mamo *et al*., 2006; Liu *et al*., 2013; Cochrane *et al*., 2015; Zhang *et al*., 2022). Larger seeds produced in warmer environments are typically attributed to longer growing seasons and increased resource availability, whereas shorter growing seasons at higher latitudes can constrain seed development (Salisbury, 1974; Stebbins, 1974; Lord *et al*., 1997; Murray *et al*., 2004). Along precipitation gradients, increased seed size has often been associated with drier environments as an adaptation that increases probability of establishment under water limited conditions (Baker, 1972; Venable, 1992; Leishman *et al*., 2000). However, these benefits reflect a fundamental trade-off between seed mass and seed number, where increased seed mass may incur a trade-off between seed number so that production of larger seeds must outweigh the potential fitness costs associated with reduced reproductive output (Smith and Fretwell, 1974; Eriksson and Jakobsson, 1999). These trade-offs extend to early life-history performance (Jones *et al*., 1997; Castro, 1999; Seiwa, 2000; Baraloto *et al*., 2005; Mamo *et al*., 2006; Warwell and Shaw, 2019; Zhang *et al*., 2022; Pawłowski *et al*., 2024). Larger and heavier seeds have been associated with increased germination, aboveground growth accumulation, and survival, particularly under stressful environments (Tripathi and Khan, 1990; Khurana and Singh, 2004; Moles and Westoby, 2004 *b*; Sánchez-Montes De Oca *et al*., 2018). These advantages can be pronounced during the transition from seed to seedling, when individuals shift from heterotrophic to autotrophic growth in more exposed environments (Donohue *et al*., 2010). Despite these benefits, they may be counterbalanced by ecological costs, including higher predation risk, greater cost associated with maternal investment, and reduced dispersal capacity (Greene and Johnson, 1993; Leishman *et al*., 2000; Gómez, 2004). As a result, characterizing these climate-dependent trade-offs is critical to understanding how life-history strategies will influence establishment and persistence. *Ex situ* seed collections serve as repositories of genetic diversity, with the potential to preserve adaptive genetic variation, support restoration, and safeguard a species’ evolutionary potential. These collections provide an invaluable resource for studying seed and seedling traits, particularly for rare or threatened species with variable seed availability (Hoban and Schlarbaum, 2014; Potter *et al*., 2017; Di Santo *et al*., 2021). As seed traits typically exhibit moderate to high heritability, variation observed within collections can serve as a proxy for quantifying genetic diversity within and among populations (Di Santo *et al*., 2021). When collections are geo-referenced, climate gradients associated with provenance origin may be associated with the evolution of ecotypic differences (Leites and Benito Garzón, 2023; Gargiulo *et al*., 2025). Quantifying trait differences within and among populations may identify geographic or environmental gaps where supplemental collections are needed (Braasch *et al*., 2021; Di Santo *et al*., 2021; Gargiulo *et al*., 2025). Thus, leveraging collections supports applied and fundamental goals associated with species conservation and restoration, informing seed sourcing strategies for restoration and management while advancing our understanding of selection and adaptation. Here, we leverage *ex situ* seed collections of critically endangered black ash (*Fraxinus nigra* Marshall) to ask how climatic gradients have influenced seed morphology and early life history traits. Specifically, we ask how climate has influenced heritable variation in these traits and how that variation affects early growth and fitness. Black ash is a foundational wetland species distributed across northeastern North America, spanning a large climatic gradient from continental to maritime environments (Springer and Dech, 2021). It also plays a central cultural and economic role for Tribal Nations in the United States and First Nations in Canada (Costanza *et al*., 2017). However, the species is severely threatened by the invasive emerald ash borer (EAB, *Agrilus planipennis* Fairmaire), which has caused substantial losses across much of the species’ range (Haack *et al*., 2002; Siegert *et al*., 2023). As a target for EAB-resistance breeding and future restoration, black ash provides a valuable system to link climatically structured variation to early life performance.

We quantified the relationship between climate of origin, seed morphology, life history traits, and first year growth using *ex situ* seed collections taken from across the range of black ash. Specifically, we (1) estimated heritability or proxies thereof for seed and early life history traits, (2) quantified associations between climate of origin and trait variation, and (3) and tested how climate of origin and variation in early life history predicted seedling fitness. This work demonstrates how *ex situ* collections can be used to identify climatically structured trait variation and guide seed sourcing, sampling, and conservation strategies aimed at maintaining adaptive trait differences under environmental change.

## Materials and Methods

### Seed Collection and Measurement

In 2023, *ex situ* seed collections of black ash were obtained from the USDA National Plant Germplasm System (NPGS) and the National Tree Seed Centre in Canada (NTSC) to evaluate variation in seed morphology and early life-history traits. Preserved seeds represent collections taken between 1991 and 2022 across the species’ native distribution. In 2023, additional seed collections were made by Pennsylvania State University and the US Forest Service to complement existing collections. In total, 76 populations containing seeds from between 3-23 georeferenced maternal lineages per population (n=701 total) were assembled from across the range of black ash (Figure 1A, Table S1). Here, we define population as a geographic location containing multiple maternal lineages with multiple seeds collected from each open-pollinated maternal lineage.

**Figure 1:**
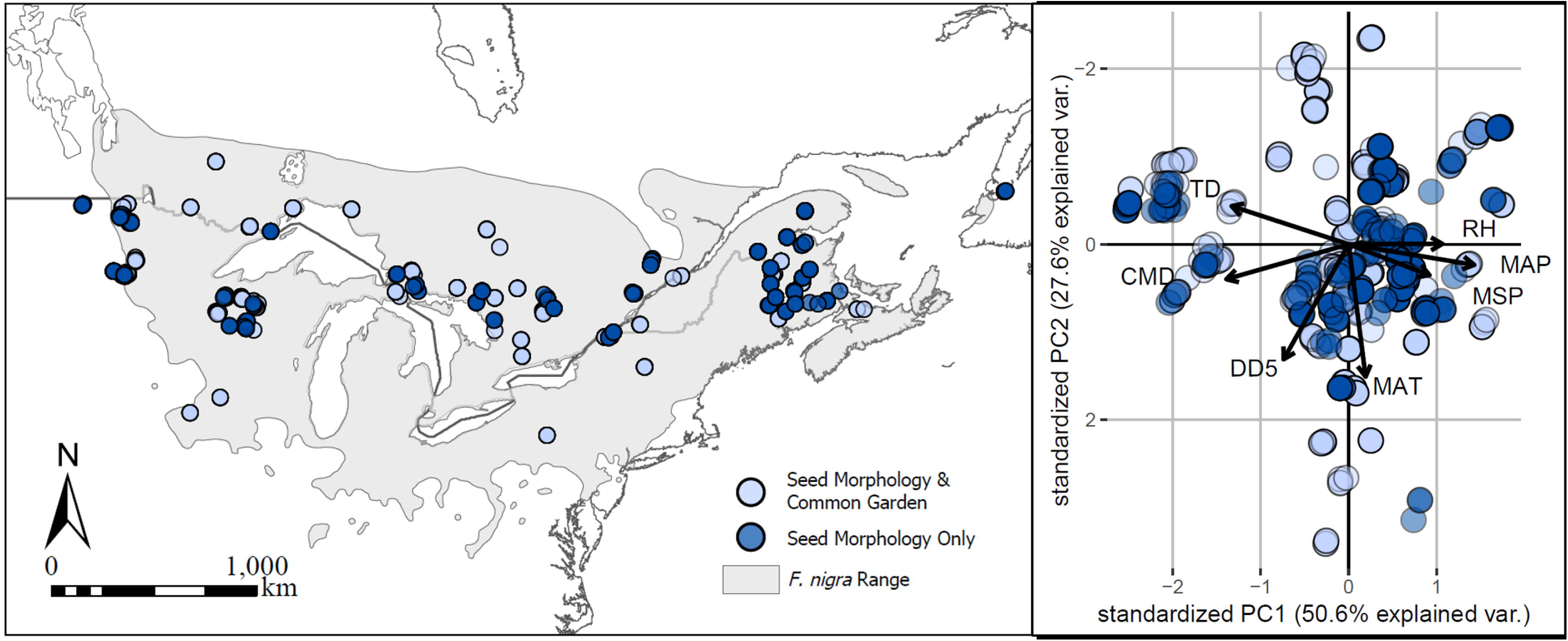
Map of ex situ seed collections for *F. nigra* (A) and principal component analysis (PCA) (B) based on seven uncorrelated climate variables associated with climate of origin obtained from ClimateNA (Wang et al., 2016). Light blue points represent maternal families used in both seed morphology and common garden experiments. Dark blue points are maternal families used for seed morphology evaluation only. The light grey polygon is the extent of the range of F. nigra based on Little’s 1971 maps. Climate variables included in the PCA are degree days greater than 5°C (DD5), mean annual temperature (MAT, °C), mean annual precipitation (MAP, mm), continentality (TD, °C), mean summer precipitation (MSP, mm), Hargreaves climate moisture deficit (CMD, mm), and relative humidity (RH, %). **Alt Text:** Map of sampling locations and principal component analysis graph of population climate data.

A random selection of 50 seeds from each of the 701 maternal lineages were x-rayed to assess morphological trait variation. X-ray images were performed at the US Forest Service National Seed Laboratory and were taken at 1x magnification on an UltraFocus digital X-ray system (Faxitron, Inc., Tuscon, AZ) at 19kV with 12 second exposures. Seeds were laid flat alongside a 10cm scale bar with no overlapping wings to ensure the entirety of the seed image was captured. Black ash seed can be divided into seed and wing structures. Seed structures include the seed coat, endosperm, and embryo, while the wing is a flat structure designed to increase surface area and facilitate wind dispersal (Figure S1).

To automate morphological measurements, an attention-gated fully convolutional neural network model (CNN) was adapted from Davaasuren *et al*., (2022) to discriminate between seed and wing structures. A subset of x-ray images was used to create a training dataset that delineated between seed, wing, and background within each x-ray image. Training images were manually segmented into seed and wing components using the ImageJ plugin ‘Labkit’ (Arzt *et al*., 2022) which produced binary mask images for seed and wing structures. All training dataset images were resized to 256 × 256 pixels for quality control and computational efficiency. To improve segmentation model robustness and generalization, flipping and rotation were applied to the training dataset. The training dataset was then divided into training (80%) and validation (20%) data. The training dataset was used to ground-truth supervised segmentation. Briefly, the CNN model performs pixel-wise classification, assigning each pixel to seed, wing, or background classes based on learned spatial and intensity features. The network architecture consists of an encoder that extracts hierarchical image features and an attention-gated decoder that emphasizes regions corresponding to target structures while suppressing background noise and imaging artifacts common in x-ray data.

The CNN model was trained using the training dataset with a maximum of 500 passes (epochs) through the data and early stopping to mitigate overfitting (Davaasuren *et al*., 2022). During each epoch, the model updated its parameters based on patterns learned from the training data, and performance was evaluated on the validation set. Convergence was assessed using the Dice coefficient (Dice, 1945), which quantifies spatial overlap between predicted segmentations and ground-truth masks. Model performance was considered to have stabilized when validation Dice scores plateaued across successive epochs, typically within ∼100 epochs. Optimization was performed using the Adam optimizer (Kingma and Ba, 2017) with a learning rate decay to promote stable convergence. Following training, the optimized CNN model was used to segment 701 x-ray images into seed, wing, and background classes. All computations were performed on an NVIDIA A100 GPU. The resulting segmented images were used for downstream analysis of seed and wing morphology. To assess segmentation accuracy, automated measurements were correlated with manual ImageJ measurements for a subset of 10 x-ray images that included 50 seeds/image (Figure S2).

Using the segmented images, the ‘centroid’ feature in ImageJ (Schindelin *et al*., 2012) was used to measure all subsequent traits for seed and wing structures.. Length (SL, WL) and width (SW, WW) measures were taken in millimeters from the longest and widest points of the seed and wing (Figure S1). Seed (SA) and wing (WA) area were calculated as square pixels occupied per structure and were converted to square millimeters using the set scale of 20.2 mm/pixel. In addition to direct morphological measures, seed to wing area ratio (SA/WA) and wing length to wing width ratio (WL/WW) were derived. Finally, for each maternal family, one hundred seeds that were not x-rayed were used to estimate total seed and wing mass (SM).

### Climate data

ClimateNA (Wang *et al*., 2016), which calculates climate averages based on a 3km grid, was used to collate historical climate data associated with population origin. Climate averages were calculated for the period 1961-1990 as this range likely reflects the historical climatic conditions that shaped the genetic composition of the sampled populations. Pairwise Pearson correlation tests were used to evaluate collinearity among climate variables. Where a correlation coefficient greater than |0.75| was observed, one variable was kept and the other was excluded (Figure S3). In total, seven climate variables were retained that both limited redundancy and were deemed ecologically relevant to the species. This included degree days greater than 5°C (DD5), mean annual temperature (MAT, °C), mean annual precipitation (MAP, mm), continentality (TD, °C), mean summer precipitation (MSP, mm), Hargreaves climate moisture deficit (CMD, mm), and relative humidity (RH, %). A scaled and centered principal component analysis (PCA) was performed using the seven climate variables to summarize axes of climate variability captured across the seed collection (Figure 1B).

### Common garden experiment and quantitative trait evaluation

A subset of populations (n=49, with between 4 to 12 maternal lineages/population) were used to establish an open-pollinated, half-sib, randomized complete block, controlled greenhouse, common garden experiment (Figure 1A). Prior to stratification, 50 non-x-rayed whole seeds from 442 maternal lineages were sterilized in 3% bleach solution and rinsed in distilled water to limit potential impacts of pests or pathogens. Sterilized seeds, grouped by maternal lineage, were then placed in a plastic bag with moist sphagnum moss and 3 inches of a 1-inch diameter clear tube running through an opening in the top of the plastic bag to ensure air exchange. Plastic bags were monitored bi-weekly to maintain moist sphagnum and monitor for potential fungal growth. As black ash exhibits morphophysiological dormancy, artificial stratification was used to facilitate maturation of the embryo and break dormancy. Plastic bags were maintained in stratification conditions between May 2023 to April 2024, starting with 90 days at 3-5°C, 90 days at 20°C, and finally 180 days at 3-5°C.

In April 2024, seeds were removed from stratification and planted under greenhouse conditions. The greenhouse was maintained between 23-29°C during the day and 12-18°C during the night at 60% relative humidity with approximately 50% shade. Individual seeds were planted in 165 well forestry trays (FT165, Stuewe and Sons, Inc.) filled with ∼90ml of a moist germination mix containing fine peat moss, perlite, vermiculite, limestone, a non-ionic wetting agent, and fertilizer (BM2, Berger). Maternal seed families within each population were randomized across benches throughout the greenhouse to minimize micro-environmental effects associated with greenhouse position. Approximately two months following emergence, seedlings were transplanted from forestry trays into 9-inch mini tree pots (MT39, Stuewe and Sons, Inc.) filled with a coarse potting mix containing coarse peat moss, coarse perlite, bark, limestone, a non-ionic wetting agent, and fertilizer (BM7, Berger). Seedlings were watered as needed throughout the course of planting and early life-history development.

Between April to September 2024, seeds were monitored for timing of seedling emergence, timing of the transition from cotyledons to true leaves, and timing of compound leaf development. Timing of seedling emergence was defined as the date when any part of the epicotyl was visible above the soil surface. The proportion of seeds that emerged given the total number of individuals planted was calculated as a measure of emergence success per maternal family. The rate at which seedlings developed cotyledons, true simple leaves, and transitioned to compound leaves were monitored daily. Finally, height measured between the base of the plant and the dormant terminal bud was recorded in September 2024 to quantify growth accumulated in the first year as a proxy for fitness. Survival was calculated for each maternal family as the proportion of seedlings alive in September 2024 relative to the number of seeds planted.

### Repeatability and heritability estimates

Direct estimation of heritability for morphological traits was not possible as seed lots were collected from different environments at different times. Given this, we calculated repeatability of traits for each population. Repeatability is defined as the proportion of trait variation attributable to differences among maternal lineages relative to the total trait variation within a population. Repeatability is often reported as an upper estimate of broad sense heritability in experimental designs where there is no way to control for environmental variance (Boake, 1989; Castro, 1999; Dohm 2002; Wolak *et al*., 2012). Our experimental design permits estimation of repeatability as trait variances were calculated for multiple maternal lineages per population and multiple individuals per maternal lineage. A linear mixed effect model was fit for each population using the R package ‘lme4’ (Bates *et al*., 2015) using R version 4.5.1 (R Core Team 2025).

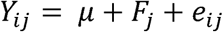

Where Y*_ijk_* is the trait of individual *i* in family *j*, µ is the overall mean, F*_j_* is the random effect of the *j*^th^ family, and e*_ij_* is the error term. Variance components were estimated using the VarCorr() function in the ‘lme4’ package (Bates *et al*., 2015) and repeatability per population was calculated as follows:

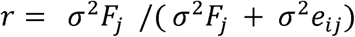

Where r is repeatability, σ^2^F*_j_* is the within family component of variance and σ^2^e*_ij_* is the error term representing residual variance not attributed to maternal families. The repeatability for each trait was calculated for each population. To estimate uncertainty in repeatability, bootstrap resampling was performed for each population. Repeatability measures were bootstrapped 1000 times by randomly resampling families with replacement to calculate repeatability variability within populations for each trait. The distribution of bootstrap values was used to calculate the 95% confidence interval and standard error around the predicted mean.

Due to the hierarchical nature of the randomized complete block, half-sib design in the greenhouse, narrow sense heritability was calculated for life history traits. To meet the assumptions of normality and homoscedasticity for linear mixed effect models, emergence timing, true leaf timing, compound leaf timing, and height were log_10_ transformed. Phenotypic variance attributed to population and maternal family were calculated using a linear mixed effect model:

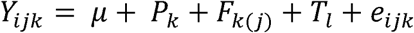

Where Y*_ijk_* is the trait of individual *i* in family *j* within population *k*, µ is the overall mean, P*_k_* is the random effect of the *k*^th^ population, F*_k(j)_* is the random effect of the *j*^th^ family within the *k*^th^ population, T*_l_* is the random effect of table, and e*_ijk_* is the error term. Table was included as a random effect to control for microsite variation within the greenhouse. Narrow sense heritability estimates use a coefficient of relatedness to account for the relationship between individuals. We used a relatedness coefficient of ¼ to account for the half sibling relationship within families since black ash is believed to be self-incompatible (Z. Smith, UT Knoxville, US, pers. comm.). Because each maternal tree was open-pollinated, some offspring may be more closely related than true half-siblings, which could lead to an overestimation of heritability. Therefore, we also estimated narrow-sense heritability using a relatedness coefficient of ½ (Table S2). From the covariances estimated in the previous equation, narrow-sense heritability was calculated as:

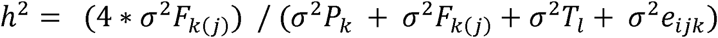

Where h^2^ is narrow sense heritability, σ^2^P*_k_* is the population component of variance, σ^2^F*_k(j)_* is the family within population component of variance, σ^2^T*_l_* is the greenhouse microsite environmental variation component of variance, and σ^2^e*_ijk_* is the error term representing within family variance. Uncertainty in heritability was quantified using a parametric bootstrap implemented with ‘bootMer’ in the R package ‘lme4’ (Bates *et al*., 2015). Specifically, 1,000 datasets were generated from the fitted mixed-effects model under an assumed Gaussian error structure, and heritability was recalculated for each generated dataset. The standard error and the 95% confidence intervals were calculated from the distribution of bootstrap heritability estimates. *Variance partitioning*

To quantify the proportion of variance attributed to population, maternal family, and individual differences for seed morphology and life history traits, we fit a linear mixed effect model:

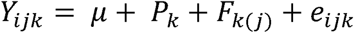

Where Y*_ijk_* is the trait of individual *i* in family *j* within population *k*, µ is the overall mean, P*_k_* is the fixed effect of the *k*^th^ population, F*_k(j)_* is the random effect of the *j*^th^ family within the *k*^th^ population, and e*_ijk_* is the error term. For each model, the significance of fixed effect terms was assessed using anova() from the R package ‘stats’ (R Core Team), and significance of random effects was assessed using ranova() from the R package ‘lmerTest’ (Kuznetsova *et al*., 2017). The proportion of variance explained by population and maternal family was quantified using the r.squaredGLMM() function in the R package ‘MuMln’ (Bartoń, 2025) to estimate the conditional R^2^ (R^2^) and marginal R^2^ (R^2^_m_). The proportion of variance explained by population was calculated as R^2^_m_, and the proportion of variance explained by maternal family was calculated as R^2^_m_ – R^2^_c_.

### Contribution of climate to trait variation

To quantify the relationship between climate of origin and seed and life history traits, linear regressions were fitted using the lm() function in R (R Core Team). Each of the seven climate variables were used to predict seed and life history traits averaged at the population level. Given no evidence of nonlinear or quadratic relationships, only linear terms were assessed. The residuals of each model were visually checked for normality and homoscedasticity to ensure the assumptions of linear regression were met.

### Relating variation in seed morphology to early life history

To evaluate the relationship between seed morphology and early life history traits, we fit linear regressions using the lm() function in R (R Core Team) between the nine seed morphological traits and five early life history traits measured in the greenhouse (proportion of seedlings emerged, timing of emergence, timing of true leaf development, timing of compound leaf development, height). The residuals of each model were visually checked for normality and homoscedasticity to ensure the assumptions of linear regression were met.

Given the correlative relationship among early life history traits and the cumulative effect of variation across multiple life stages to fitness, we also used an aster life history analysis to describe how traits cumulatively effect fitness over time (Geyer *et al*., 2007; Shaw *et al*., 2008). Aster analysis jointly models multiple components of fitness with different probability distributions, accounting for the dependence of variables on earlier stages to estimate fitness as an integrated measure of life history across an individual’s lifespan (Geyer *et al*., 2007). Each life history component is represented as a node in a graphical model with the distribution of each node conditioned on the node preceding it. In this way, the graphical model becomes the response matrix and fixed and random effects are added as predictors to estimate the effect of each trait of interest to fitness. To quantify the effect of early life history trait variation to proxies of first year fitness for a perennial tree species, we included seed mass, timing of seedling emergence, true leaf development, compound leaf development, and the presence/absence of true and compound leaves as fixed effects. Population was also included as a fixed effect and maternal family within population was treated as a random effect. To estimate fitness following the first year, we included survival modelled as a Bernoulli distribution and first year height as a Gaussian distribution in the graphical model. We could not assess lifetime fitness for seedlings given their long generation time. However, as height has been shown to correlate with greater reproductive output in forest trees (Klinkhamer *et al*., 1997), we consider accumulation of height in the first year of growth conditioned on survival to be the terminal fitness node in our model. A conceptual figure showing the graphical model and how each life history component fits into the model can be found in Figure S4.

### Contribution of climate to cumulative fitness

In addition to quantifying the effects of life history to fitness following first year of growth, the aster model was used to predict population-level fitness. Following Kulbaba *et al*. (2019), fitness predictions were obtained by transforming model estimates from the canonical (link) scale to the mean-value scale. Specifically, fixed effects were combined with each family-level random effect on the canonical scale to generate family-specific linear predictors. These values were then back-transformed to the mean-value scale using the inverse link (mapping) function, yielding expected fitness for each family. Population-level fitness (µ), survival, and height were calculated by averaging these family-level fitness estimates. Linear regression was used to test if climate of origin predicted mean predicted population fitness following first year of growth. Residuals of each model were visually checked for normality and homoscedasticity to ensure the assumptions of linear regression were met.

## Results

### Repeatability of seed morphology traits and heritability of early life history traits

Here we report repeatability as a proxy for broad-sense heritability for seed morphological traits. On average, repeatability for seed traits was 0.41 (range = 0.30-0.54), with wing length to width ratio exhibiting the greatest repeatability (0.54) and seed to wing area ratio exhibiting the lowest (0.30) (Table 1). The bootstrapped mean repeatability was slightly lower than the observed mean for all traits, which may be attributed to the right skew of the data (Figure S5). However, the 95% confidence intervals were narrow, indicating a predictable central tendency for repeatability. Mean repeatability values for each trait per population can be found in Table S3. Narrow sense heritability was measured for life history traits in the greenhouse common garden experiment. The heritability of traits ranged from 0.83 ± 0.06 for emergence timing, 0.39 ± 0.04 for the timing of true leaf development, 0.49 ± 0.08 for the transition to compound leaves, and 0.83 ± 0.07 for height (Table 1). In addition, additive genetic variance was measured (V_A_) to provide an estimate of genetic variation within traits available for selection to act upon. Compound leaf development had the greatest additive genetic variance (0.127), followed by emergence (0.069), height (0.064), and true leaf development (0.018, Table 1). These heritability estimates are likely slightly inflated due to the log_10_ transformation of raw data. As the transformation compresses higher values, the resulting heritability estimates reflect the proportion of variance on a relative (log) scale, which can increase h² compared with estimates based on untransformed data.

**Table 1:**
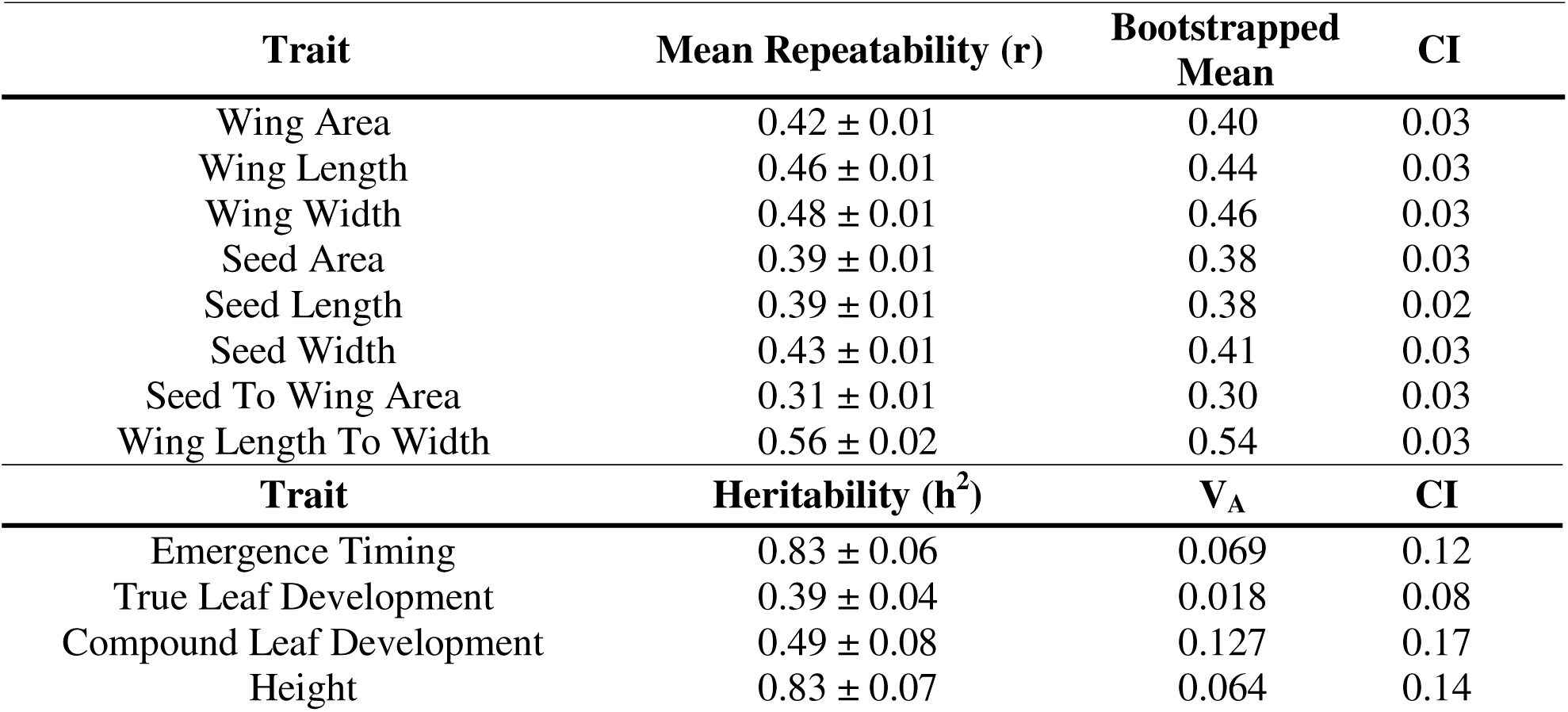
Repeatability estimates measured as a proxy for broad-sense heritability for seed traits averaged across populations and narrow sense heritability estimates for life history traits measured under controlled environment conditions in a common garden experiment. A distribution of repeatability measures was calculated using a bootstrap with replacement approach within each population for 1,000 replicates. For narrow-sense heritability, a distribution of heritability measures was calculated using a parametric bootstrap approach with 1,000 replicates per trait. 95% confidence intervals (CI) were derived from the bootstrap distribution for both seed morphology and life history traits. For both repeatability and heritability estimates, the standard error is reported after the mean. For early life history traits, additive genetic variance is also reported (V_A_).

| Trait | Mean Repeatability (r) | Bootstrapped Mean | CI |
| --- | --- | --- | --- |
| Wing Area | 0.42 ± 0.01 | 0.40 | 0.03 |
| Wing Length | 0.46 ± 0.01 | 0.44 | 0.03 |
| Wing Width | 0.48 ± 0.01 | 0.46 | 0.03 |
| Seed Area | 0.39 ± 0.01 | 0.38 | 0.03 |
| Seed Length | 0.39 ± 0.01 | 0.38 | 0.02 |
| Seed Width | 0.43 ± 0.01 | 0.41 | 0.03 |
| Seed To Wing Area | 0.31 ± 0.01 | 0.30 | 0.03 |
| Wing Length To Width | 0.56 ± 0.02 | 0.54 | 0.03 |
| **Trait** | **Heritability (h<sup>2</sup>)** | **V<sub>A</sub>** | **CI** |
| Emergence Timing | 0.83 ± 0.06 | 0.069 | 0.12 |
| True Leaf Development | 0.39 ± 0.04 | 0.018 | 0.08 |
| Compound Leaf Development | 0.49 ± 0.08 | 0.127 | 0.17 |
| Height | 0.83 ± 0.07 | 0.064 | 0.14 |

### Within and among population trait differentiation

Both population and maternal family contributed substantially to differences observed across traits (Figure 2, Table S4). On average, population explained ∼14% (range = 0.09-0.17) of the morphological trait differences observed given the populations assessed. Population explained the greatest variance in the wing traits, including wing area (0.17), wing width (0.17), and wing length (0.15). For early life history traits, population differences explained ∼4.5% of the variation in timing of emergence and leaf development, and ∼11% of the among-population differences in height (Table S4). Population had no significant impact on the timing of compound leaf development.

**Figure 2:**
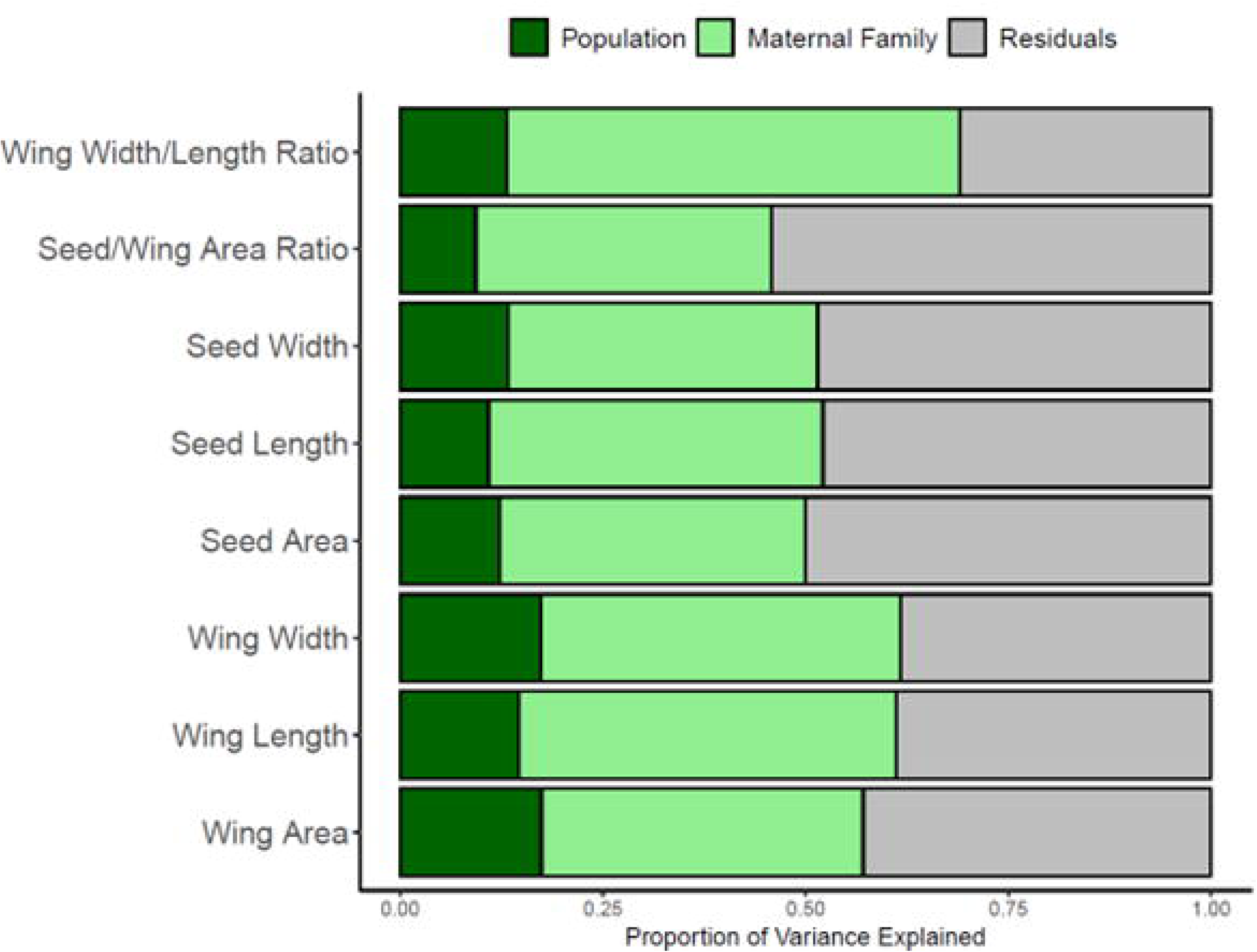
Proportion of variance in seed morphological traits explained by populations (dark green), maternal families within populations (light green), and other variables not included in the model (grey) for each of the 8 measured and derived traits. **Alt Text:** Bar graph of the proportion of variance explained by population, maternal family, and residuals.

On average, family explained 43% (range 0.37-0.56) of the observed differences in seed morphology (Figure 2, Table S4). The highest proportion of variance attributed to maternal family was for wing width to length ratio (0.56), wing length (0.47), and wing width (0.44). Among early life history traits, maternal family explained 21% of the variability in timing of emergence, 10% in timing of true leaf development, 15% for the transition to compound leaves, and 23% for height accumulated in the first year (Table S4).

### Seed morphology and life history trait variation associated with climate of origin

A PCA based on climate of population origin indicated that populations were primarily structured by variation in temperature and precipitation. PC1 explained 50.6% of the variation observed and was primarily influenced by precipitation related variables (MAP, MSP, CMD), and PC2 explained 27.6 % of the variability observed, with loadings primarily temperature-related variables (MAT, DD5) (Figure 1B, Table S5).

Linear regressions of climatic with morphology and life history traits indicate that mean annual temperature (MAT), growing degree days over 5°C (DD5), mean summer precipitation (MSP), and climate moisture deficit (CMD) predict trait variability (Figure 3, Table S6). Warmer and drier environments were associated with larger and heavier seeds and increased accumulation of first year height. Further, seedlings from warmer environments displayed faster developmental transitions. Along precipitation gradients, trees from environments with greater precipitation on average produced smaller seeds and emerged more rapidly.

**Figure 3:**
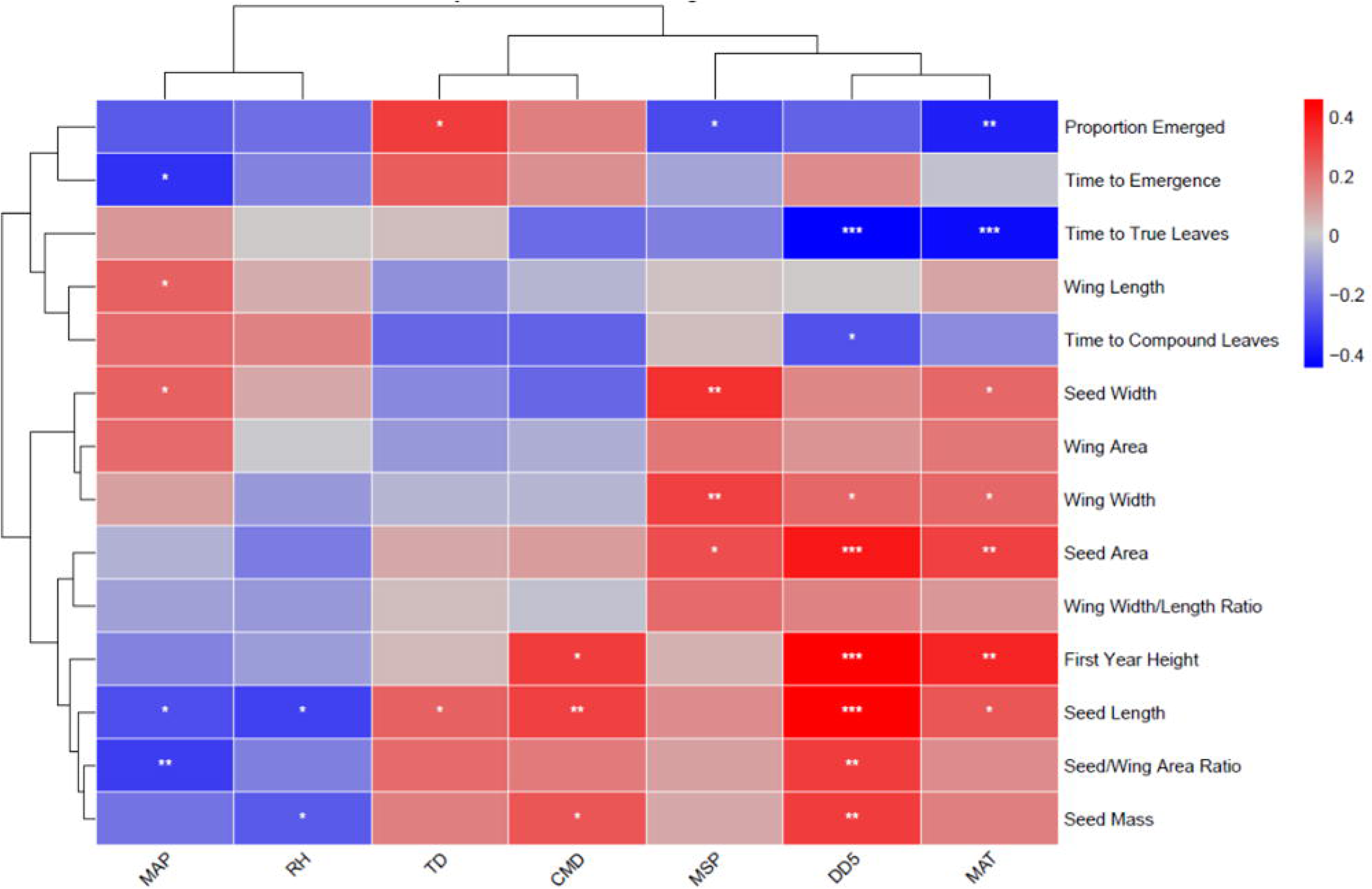
Heatmap of linear regression estimates between climate variables and all seed morphology and life history traits. Estimates were standardized to allow comparisons between traits with the strength of relationships. Each square is filled based on standardized estimates, with red values indicating a positive slope, and blue values indicating a negative slope. Stars represent the significance level. Climate variables tested are degree days greater than 5°C (DD5), mean annual temperature (MAT, °C), mean annual precipitation (MAP, mm), continentality (TD, °C), mean summer precipitation (MSP, mm), Hargreaves climate moisture deficit (CMD, mm), and relative humidity (RH, %). * p < 0.05 ** p < 0.01 ***p < 0.001 **Alt Text:** Heatmap showing standardized linear regression coefficients between climate variables and seed morphology and life-history traits.

### Seed morphology predicts early life history success

On average, seed mass was the best predictor of seedling emergence (β = 0.08/mg, R^2^ = 0.10), the time to transition between cotyledons and true leaves (β = −0.42 days/mg, R^2^ = 0.36), and the accumulation of first year height (β = 0.60cm/mg, R^2^ = 0.37) (Figure 4, Table S7). Increased seed mass, seed and wing length, seed and wing width, and seed and wing area were associated with faster timing of true leaf development (Table S7, Figure S6). Despite these relationships, seed morphology did not predict variance in emergence or developmental transitions from simple to compound leaves. Furthermore, wing length to width ratio and seed to wing area ratio exhibited no significant relationship with early life history traits.

**Figure 4:**
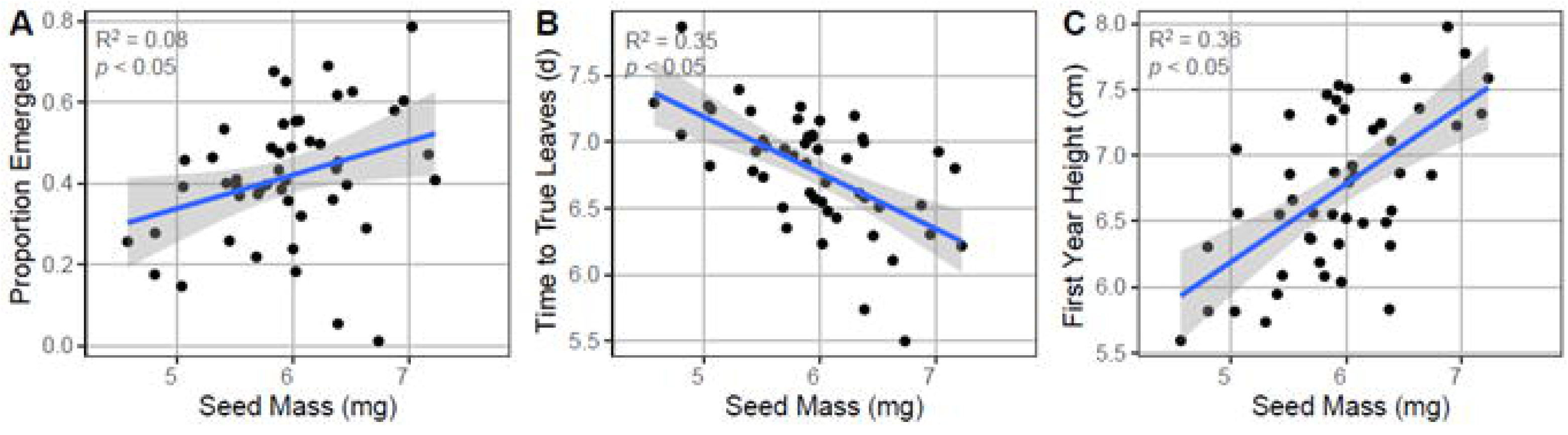
Linear regression between seed mass (n=100) and (A) proportion of seedlings emerged, (B) time to true leaf emergence, and (C) first year height. Each point represents the mean value for one population. The solid blue line represents the fitted linear regression, and the shaded area is the 95% confidence interval. **Alt Text:** Graphs showing the linear regressions between seed mass and proportion of seedlings emerged, time to true leaf emergence, and first year height.

### Population, seed mass, and timing of life history transitions predict cumulative fitness

Aster life history models indicated that population, seed mass, and all early life-history traits had a significant effect on the first-year height conditioned on survival, used as a proxy for fitness (Table 2). Seed weight, timing of seedling emergence, true leaf presence, and compound leaf presence all had a positive and significant effect on fitness at the end of the first year of growth. Heavier seeds, and seedlings that produced true leaves and compound leaves had higher survival probability and accumulated greater height in the first year on average. The timing of seedling emergence exhibited a slight but significantly positive relationship with cumulative fitness (β = 0.04 mu/day), indicating that seedlings that emerged later had greater aboveground growth accumulated in the first year. The timing of true leaf and compound leaf development had negative relationships with cumulative fitness, suggesting that seedlings that took longer to make these early life history transitions have reduced survival or reduced overall aboveground growth in the first year.

**Table 2:**
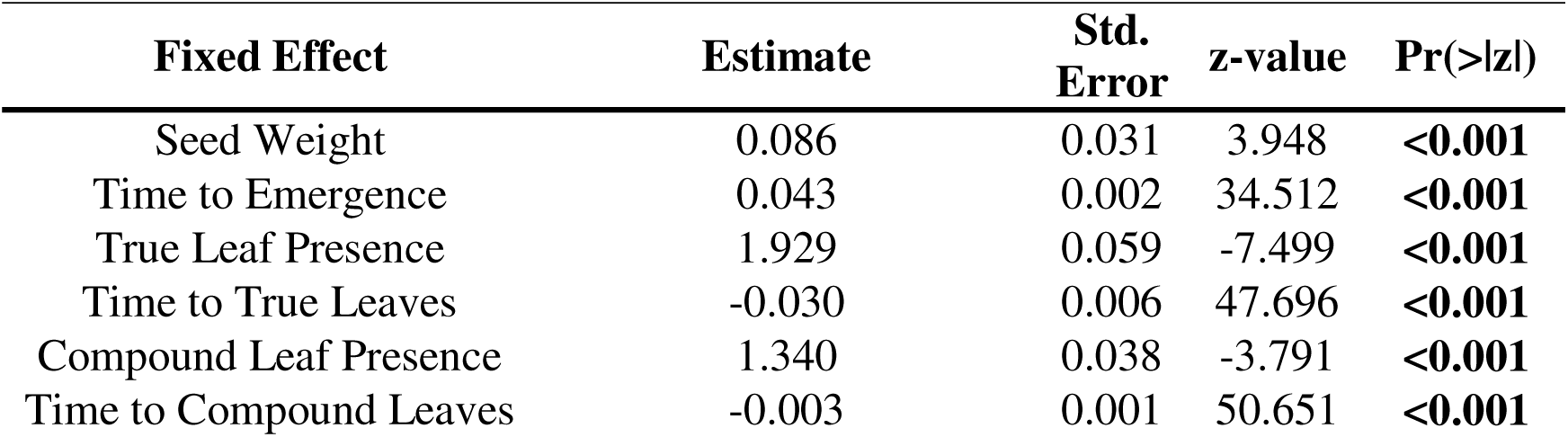
Results for fixed effect estimates for seed weight and early life history traits from the aster life history model. Fixed effects were estimated via maximum likelihood and standard errors were obtained from the inverse observed Fisher information matrix. Estimates were converted from canonical values to marginal mean estimates. Significance was assessed using Wald z-tests. Significant p-values are bolded.

| **Fixed Effect** | **Estimate** | **Std. Error** | **z-value** | **Pr(>\|z\|)** |
| --- | --- | --- | --- | --- |
| Seed Weight | 0.086 | 0.031 | 3.948 | **<0.001** |
| Time to Emergence | 0.043 | 0.002 | 34.512 | **<0.001** |
| True Leaf Presence | 1.929 | 0.059 | -7.499 | **<0.001** |
| Time to True Leaves | -0.030 | 0.006 | 47.696 | **<0.001** |
| Compound Leaf Presence | 1.340 | 0.038 | -3.791 | **<0.001** |
| Time to Compound Leaves | -0.003 | 0.001 | 50.651 | **<0.001** |

### First year height is explained by climate of origin

We tested if climate of origin predicted fitness in the first year of growth as estimated from aster life history models. We found that continentality (°C) was the best predictor of cumulative fitness (β = 0.13 µ/°C, R^2^ = 0.10), with seedlings from more continental environments having greater first year projected fitness (Figure 5, Table S8). The relationships between climate of origin and the major fitness nodes, height and survival, indicated that greater climate moisture deficit predicted greater height in the first year (β = 0.01 cm/mm, R^2^ = 0.09), and increased continentality and reduced mean annual temperature predict increased survival (β = 0.11/°C, R^2^ = 0.10) (β = −0.04/°C, R^2^ = 0.13) (Table S8).

**Figure 5:**
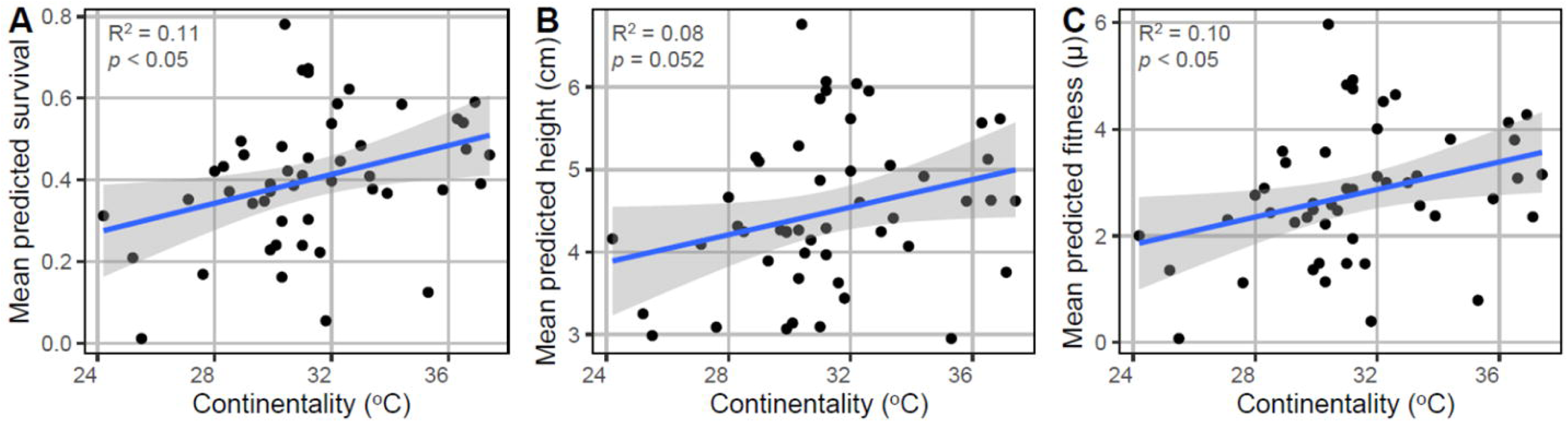
Linear regressions between continentality of the population origin and mean predicted survival (A), height (B), and fitness (C) for each population. Each point represents the mean value for one population. The solid line represents the fitted linear regression, and the shaded area is the 95% confidence interval. **Alt Text:** Graphs showing the linear regressions between continentality and survival, height, and fitness.

## Discussion

Climate gradients can select for seed morphology and life history traits that cumulatively impact establishment and early life fitness in perennial systems (Leishman *et al*., 2000; Donohue *et al*., 2010). Here, we identified heritable variation in seed morphology and early life history traits within and among black ash populations associated with climate gradients. Warmer and drier climates were associated with larger, heavier seeds, faster developmental transitions, and greater accumulation of seedling height. These differences suggest climatically structured seed trait variation can have cascading effects on establishment. Populations from more continental environments, characterized by greater seasonal temperature variability and higher climate moisture deficits, tended to produce heavier seeds and exhibited higher cumulative fitness in the first year. These patterns may reflect the evolution of trade-offs for maternal investment needed in more uncertain environments to ensure early establishment in lieu of dispersal potential (Smith and Fretwell, 1974; Venable, 1992; Eriksson and Jakobsson, 1999). As temperature variability and extreme events are projected to increase across much of the species’ range, populations from more continental environments may possess traits that buffer seedling establishment during climatic uncertainty. These findings demonstrate that climate of origin shapes seed traits and developmental timing in ways that directly influence early-life fitness. For *ex situ* conservation, this highlights the importance of collecting both within- and among-population variation across the climatic distribution of black ash. Maintaining diversity will be particularly urgent given ongoing population declines due to emerald ash borer, as it will be needed to ensure evolutionary potential is preserved to respond to shifting conditions and balance key life-history trade-offs influencing establishment.

### Within and between population variation for seed morphology is influenced by climate gradients

Repeatable differences across seed traits suggest seed morphology provides a reasonable proxy to quantify heritable within-population genetic variation (Roy *et al.,* 2004; Carles *et al*., 2009; Zas and Sampedro, 2015; Di Santo *et al*., 2021). In addition, narrow sense heritability estimates for life history traits suggest these traits will respond to natural selection, particularly emergence timing and height, which exhibit a substantial amount of additive genetic variance (Table 1). Overall, within-population variation observed across traits indicates that black ash populations retain substantial standing genetic variation upon which selection may act (Table 1, Figure 2). However, it is also clear that among population differences have evolved, suggesting potential evolution of ecotypic differences (Figure 2, Table S4). The amount of within population variation observed for traits underscores the importance of sampling multiple maternal lineages within populations to ensure standing genetic variation is captured within *ex situ* collections (Hoban and Schlarbaum, 2014; Di Santo and Hamilton, 2021; Di Santo *et al*., 2021; O’Brien *et al*., 2025). Maintaining genetic diversity within and among populations will be particularly important under climate change, ensuring populations have the capacity to adapt to change.

### Climate-mediated seed trait allocation drives variation in developmental timing and early fitness

Longer growing seasons and greater carbon accumulation may underly the positive relationship between temperature and seed size observed here, consistent with patterns observed across taxa (Figure 3, Table S6) (Moles and Westoby, 2003; Moles *et al*., 2004, 2005; Soper Gorden *et al*., 2016; Lenzo *et al*., 2024). However, the negative relationship observed between moisture and seed size is less consistent across species (Figure 3, Table S6) (Moles *et al*., 2005; Mamo *et al*., 2006; Soper Gorden *et al*., 2016; Zhang *et al*., 2022). In black ash, our results point to a shift in allocation between dispersal and seed provisioning structures along climate gradients. As a wind-dispersed diaspore, black ash faces a fundamental trade-off between investment in dispersal structures that enhance dispersal potential and seed tissues that support establishment (Greene and Johnson, 1993). Increased seed mass in warmer, drier environments suggests selection for seed structure provisioning where moisture may be limiting to early survival. Under these conditions, investment in seed provisioning may buffer environmental stress and ensure faster developmental transitions needed to outcompete neighbors for limited resources available (Leishman *et al*., 2000; Moles and Westoby, 2004 *a*,*b*) (Figure 4). Conversely, in cooler or wetter environments where establishment uncertainty is reduced, selection may favor investment in dispersal structures, promoting spread to limit local competition. This suggests that black ash may have evolved in response to varying environmental uncertainty, with investment in seed structures following resource availability gradients needed to persist under different climatic conditions.

Because fitness can reflect the cumulative impact of variation across an individual’s life history, aster models were used to evaluate which components of an individual’s early life history most impact fitness in the first year of growth (Shaw *et al*., 2008; Wadgymar *et al*., 2024). Seed mass, developmental timing, and production of leaves jointly affect first year growth, measured here as height conditioned on survival (Table 2). Seedlings with heavier seeds and faster leaf development exhibited higher cumulative fitness. Larger seed size likely reflects evolved differences in maternal investment associated with trade-offs between dispersal and establishment. This increased provisioning buffers offspring with nutrient reserves, potentially allowing delayed germination under uncertain conditions while maintaining establishment potential (Leishman *et al*., 2000; Seiwa, 2000; Moles and Westoby, 2004). Interestingly, slightly delayed emergence was associated with increased fitness. This suggests that emergence timing and subsequent developmental rate may represent different axes of an early life history strategy. Delayed emergence may enable increased allocation to belowground development prior to shoot emergence, enhancing early resource acquisition and establishment (De La Fuente Cantó *et al*., 2020). Once emerged, rapid transitions from cotyledons to true and compound leaves likely accelerate both carbon sequestration aboveground and nutrient acquisition belowground, improving competitive ability in the shaded environments where black ash seedlings typically establish (Reich *et al*., 1998, Kidson and Westoby, 2000; Moles and Westoby, 2004 *b*). However, accelerated leaf development may increase exposure to frost damage or drought stress (Seidel and Menzel, 2016; Baumgarten *et al*., 2023). Cumulatively, these patterns suggest that early establishment in black ash reflects the evolution of a coordinated life-history strategy that links maternal investment with developmental timing and resource acquisition.

### Populations from more continental environments have greater survival and cumulative growth

Climate of origin predicted cumulative aboveground growth, a proxy for fitness in the first year as estimated using the aster life history model, with continentality being the best predictor of both first-year height and survival (Figure 5, Table S8). Continental environments are characterized by greater seasonal temperature extremes, reduced precipitation, and increased uncertainty during seasonal transitions compared to more moderate coastal environments. Such environments can select for life-history strategies that buffer environmental uncertainty, including increased maternal provisioning, bet-hedging in germination timing, and increased plasticity (Anderson *et al*., 2012; Vázquez *et al*., 2017; Stotz *et al*., 2021; Volk *et al*., 2022). Within this context, increased growth accumulated in black ash populations from more continental regions may reflect the evolution of life history strategies that enhance establishment under uncertain environments. Traits that promote reliable establishment despite climatic extremes, such as heavier and larger seeds, suggest the evolution of greater maternal provisioning under uncertain environments. Theoretical expectations suggest that increased investment per offspring is favored in unpredictable or resource-limited environments because it enhances the probability of establishment under adverse conditions (Venable, 1992; Leishman *et al*., 2000; Moles and Westoby, 2004 *b*). In this way, increased seed mass may function as a buffering mechanism, allowing seedlings to tolerate the variability of continental climates. With climate change, increased temperature variability and extreme events are projected across much of the current black ash range (U.S. NOAA Climate Explorer, 2025). Populations originating from more continental environments may therefore possess traits that confer resilience to such variability, including greater provisioning and coordinated developmental strategies that enhance both survival and early growth under fluctuating conditions.

As maternal investments were not controlled for within our experiment, seed and wing trait development could reflect the impact of the maternal growing environment (Galloway, 2005; Donohue, 2009). While this complicates interpretation of population-level adaptation, maternal provisioning itself is an ecologically meaningful mechanism through which plants influence offspring establishment under local environmental conditions. Because the relationships between climate, seed traits, and seedling performance are consistent across populations they likely reflect both maternal provisioning effects and the evolution of adaptive genetic differences. Monitoring seedlings longer term will be needed to disentangle maternal environmental effects from genetic adaptation, as we predict that the potential impact of the maternal effect on progeny trait variation may diminish over time.

### Implications for conservation and management

Variation in seed morphology and early life-history traits within black ash populations indicates the species likely retains considerable standing genetic variation required to maintain evolutionary potential. However, the variation identified between populations suggests that genetic differences have evolved across the species’ range, reflecting diverse morphological and life history traits shaped by natural selection. Thus, effective conservation of black ash will rely upon preservation of both within population variation to preserve the species’ evolutionary potential and among population genetic differences to maintain ecotypic differences across the species’ range. For *ex situ* conservation efforts, this implies sampling should be structured to capture variation at both levels. Within populations, seeds should be collected from 25-30 maternal trees, following recommendations made by Hoban and Schlarbaum (2021) to capture 95% of genetic variation. Across populations, collections should span the species’ climatic niche to ensure representation of environmentally structured genetic differences (Hoban and Schlarbaum, 2014; Di Santo *et al*., 2021). This will be particularly urgent given ongoing population declines due to the emerald ash borer. Preservation of these seed ensures that there is a foundation laid for future efforts associated with breeding, restoration, or resistance screening (McManus *et al*., 2025).

## Conclusion

Here, we show that climate has acted as a selective force shaping heritable variation in seed morphology and early life-history traits for black ash. Genetic variation observed within and among populations for seed and life history traits suggests that there is substantial standing genetic variation maintained in *ex situ* collections that may be important to species adaptation. Populations originating from warmer, drier, and more continental environments produced larger seeds and developed more rapidly through early life history stages, which translated into greater first year growth. These patterns align with ecological theory that predicts maternal investment enhances establishment, particularly under resource-limited or variable environments, reflecting trade-offs between local establishment and dispersal under uncertain conditions. The climatically structured variation observed across populations underscores the importance of conserving both within- and among-population diversity for black ash. *Ex situ* seed collections that capture within and among population diversity provide a critical mechanism for preserving adaptive variation and supporting restoration, breeding, and assisted regeneration. In the context of ongoing population decline due to the emerald ash borer and increasing climatic variability, ensuring conservation targets within- and among-population diversity will be critical to future restoration efforts.

## Supporting Information

The following additional information is available in the supplemental material:

Table S1. Metadata for all assembled populations.

Table S2. Comparison of narrow sense heritability estimates for greenhouse traits using a relatedness coefficient of 1/2 and 1/4.

Table S3. Repeatability estimates for each population for each seed morphology trait.

Table S4. Variance partitioning at the population, family, and individual level for all seed morphology and life history traits. Variance components were estimated from mixed-effect models. P-values for fixed effects were calculated using ANOVA F-tests.

Table S5. Principle component (PC) loadings for the seven selected climate variables obtained from ClimateNA for each population.

Table S6. Slope estimates from linear regressions of climate variables with seed morphological traits and early life history traits. For linear regression, all traits were analyzed at the population mean level. Significance of regression coefficients was assessed using two-sided t-tests.

Table S7. Slope estimates from linear regressions of seed morphological traits with early life history traits. Significance of regression coefficients was assessed using two-sided t-tests.

Table S8. Slope estimates from linear regressions of historic climate variables with overall fitness and fitness components from the aster models averaged per population. Significance of regression coefficients was assessed using two-sided t-tests.

Figure S1. X-ray image of seeds with morphological traits measured indicated. Wing length (WL), wing width (WW), and wing area (WA) measure the entire surface of the samara, while seed length (SL), seed width (SW), and seed area (SA) measure the seed only.

Figure S2. Correlations between manual and automatic measurements of wing length (A), wing width (B), seed length (C), and seed width (D) for a subset of ten maternal lineages.

Figure S3. Pearson correlation coefficients between all 25 annual average climate variables from 1961-1990 from ClimateNA (Wang *et al*., 2016).

Figure S4. Conceptual figure showing the components of the model used for aster analysis.

Figure S5. Histograms and qq-plots to visualize normality of the 9 measured and derived seed morphology traits.

Figure S6. Pearson correlation coefficients between seed morphological traits and early life history traits.

## Availability of Data and Materials

All data and scripts associated with this manuscript are available on GitHub (https://github.com/krlopicc/SeedMorphology_LifeHistory). Details on the CNN model can be found at https://github.com/Dolzodmaa/GuardCellSegmentation.

## Author Contributions

KL led data collection, research design, data analysis and interpretation, and manuscript writing. JM contributed to data collection, data analysis and interpretation, and manuscript writing. LA and DD contributed to data collection and manuscript writing. JAH contributed to data collection, research design, data analysis and interpretation, and manuscript writing.

## Acknowledgments

We thank Jeff Carstens at USDA-ARS National Plant Germplasm System and Donnie McPhee at the National Tree Seed Centre for supplying black ash seeds from existing *ex situ* collections. We also thank Nate Siegert at the US Forest Service, Maureen Beeman, and Nadia Garzione for contemporary seed collections. We’re grateful to Tia Tyler and Victor Vankus at the US Forest Service National Seed Laboratory for taking the x-ray images. Additional thanks go to Melissa Lehrer for her contributions to organizing seed collections and stratification, to Alex Moen for assistance with data collection, and to Sam Gruneberg for help maintaining seedlings in the greenhouse. Finally, we appreciate Michelle Zavala-Paez, Alayna Mead, Diego del Orbe, Mary McCafferty, Muqing Liu, and Sammy Muraguri from the Hamilton lab for their questions, discussion, and feedback during lab meetings, which helped shape the direction of this work.

## AI Assistance Acknowledgement

AI tools (specifically ChatGPT) were used only to assist with rephrasing text and formatting code. All scientific content, analysis, and conclusions were generated and verified by the authors.

