## Supplemental Figures for "Climate gradients drive the evolution of seed morphology and life history with impacts to seedling fitness in *Fraxinus nigra*"

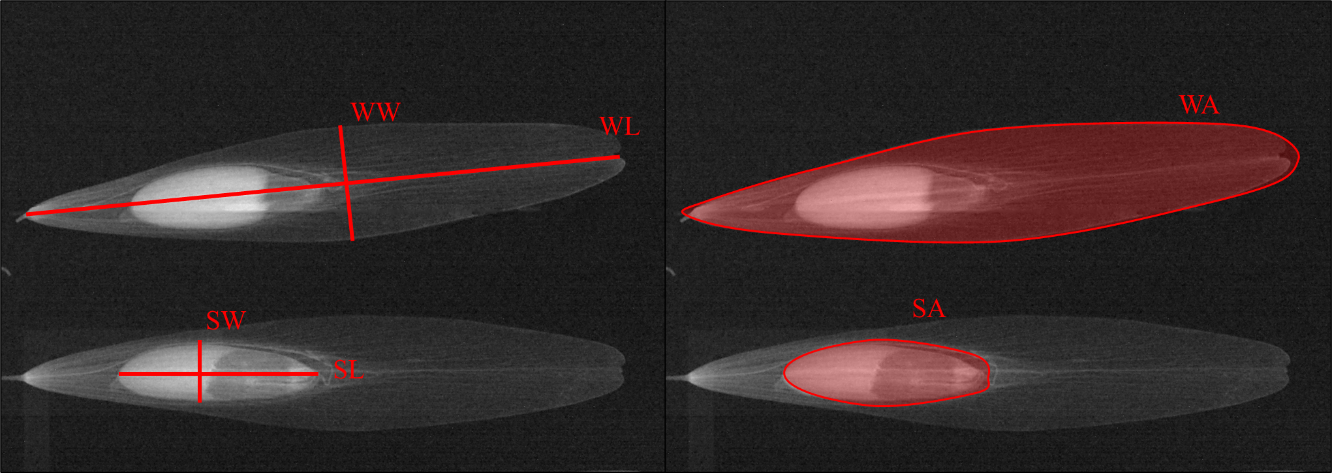
 **Supplementary Figure 1:** X-ray image of seeds with morphological traits measured indicated. Wing length (WL), wing width (WW), and wing area (WA) measure the entire surface of the samara, while seed length (SL), seed width (SW), and seed area (SA) measure the seed only.

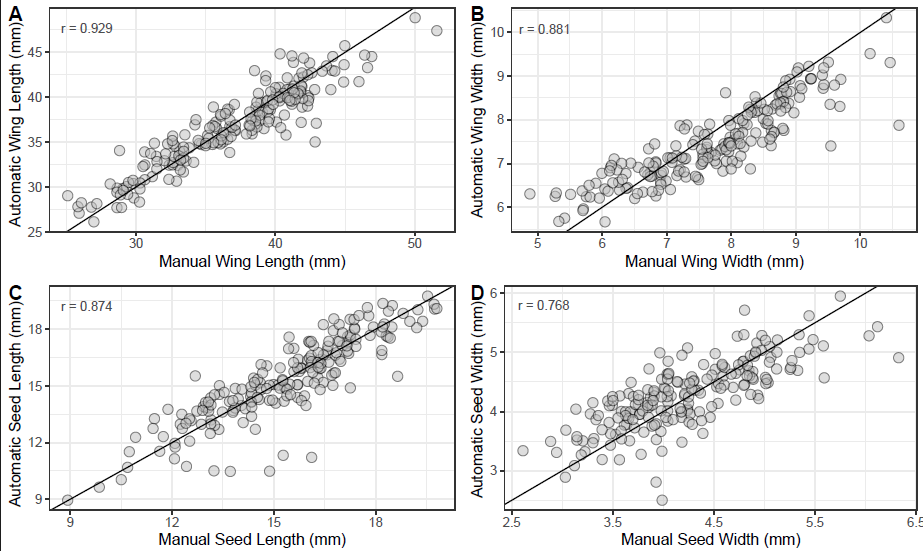

**Supplementary Figure 2:** Correlations between manual and automatic measurements of wing length (A), wing width (B), seed length (C), and seed width (D) for a subset of ten maternal lineages. Pearson correlation coefficients are displayed in the top left corner. The solid black line represents the 1:1 relationship.

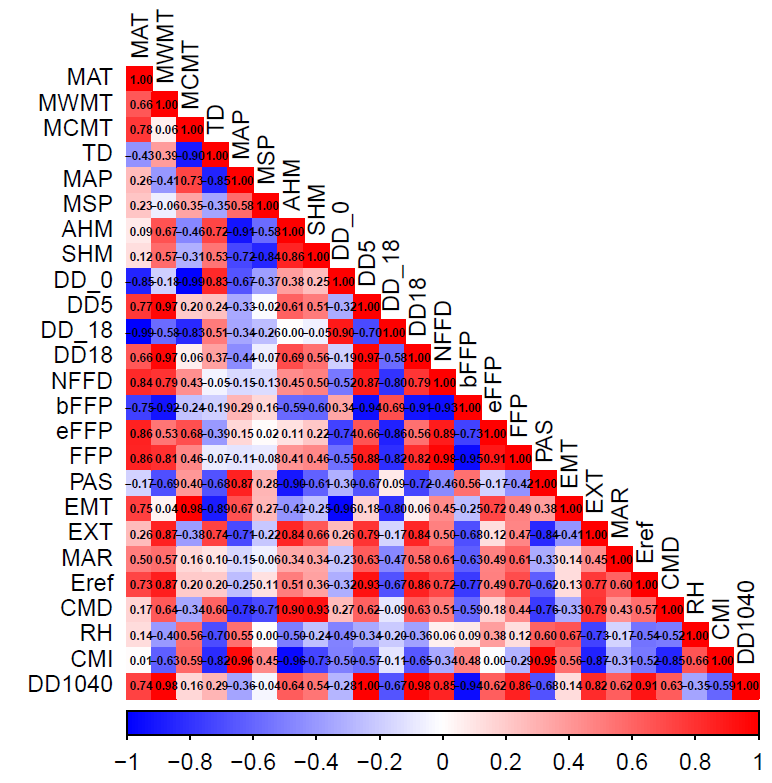

**Supplementary Figure 3:** Pearson correlation coefficients between all 25 annual average climate variables from 1961-1990 from ClimateNA (Wang et al., 2016).

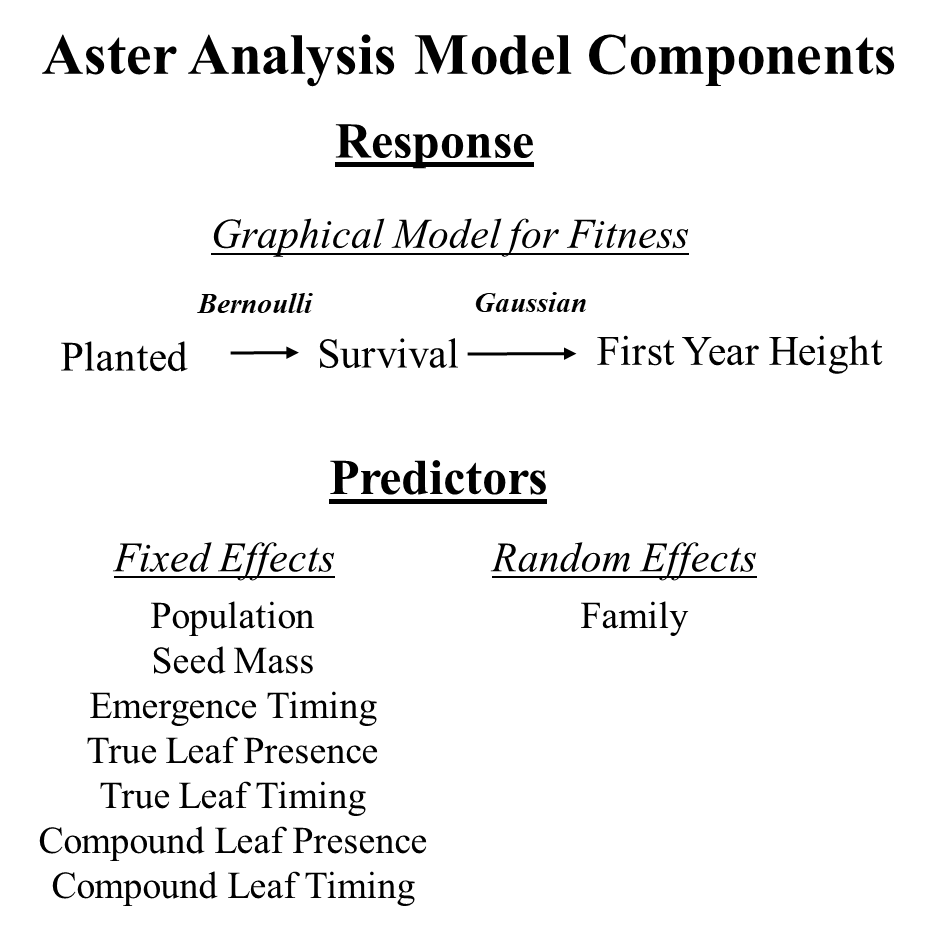

**Supplementary Figure 4:** Conceptual figure showing the components of the model used for aster analysis. The response variable in the model is a graphical model of fitness where each node represents a component of fitness. The cumulative value at the terminal fitness node, in this case height conditioned on survival, is the overall fitness. The predictors are variables of interest that we want to quantify the relationship of with overall fitness.

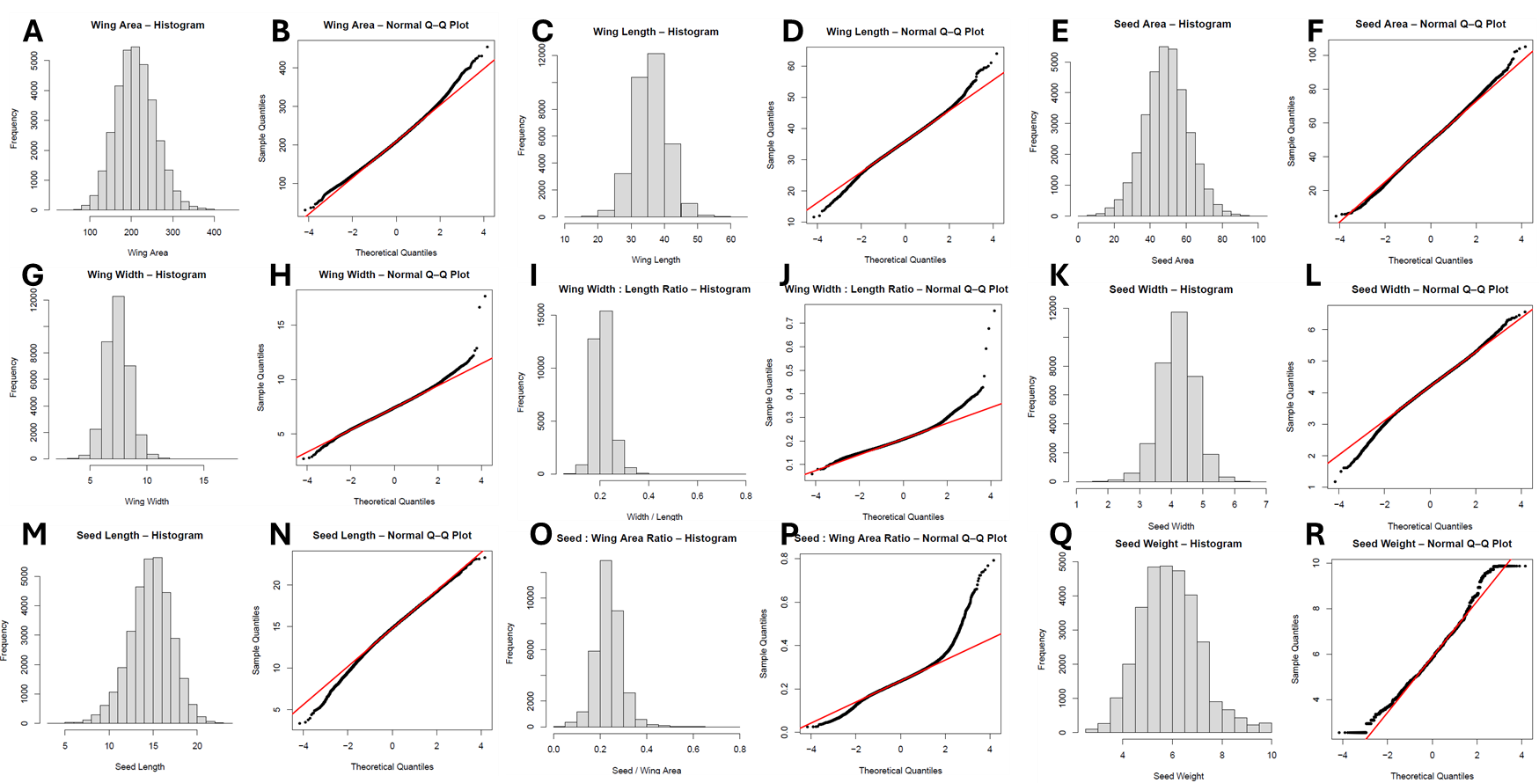
**Supplementary Figure 5:** Histograms and qq-plots to visualize normality of the 9 measured and derived seed morphology traits.

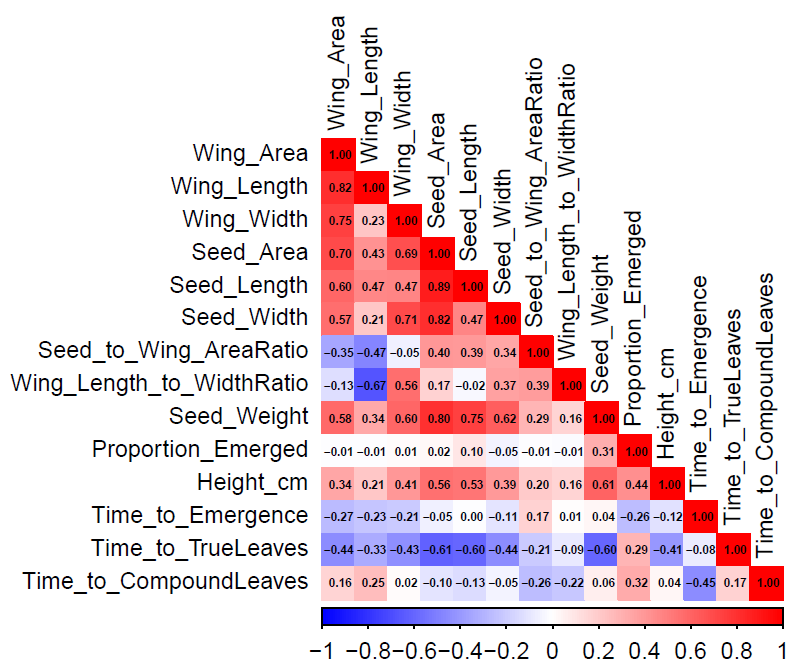

**Supplementary Figure 6:** Pearson correlation coefficients between seed morphological traits and early life history traits**.**
